# Quinoclamine inhibits Shiga toxin production in enterohemorrhagic *Escherichia coli*

**DOI:** 10.1101/2023.09.26.559460

**Authors:** Oiti Kar, Hsiao-Cheng Feng, Hiroyuki Hirano, Ching-Hao Teng, Hiroyuki Osada, Masayuki Hashimoto

**Affiliations:** Institute of Molecular Medicine, College of Medicine, National Cheng Kung University, Tainan, 701, Taiwan; Chemical Resource Development Research Unit, Technology Platform Division, RIKEN Center for Sustainable Resource Science, Saitama 351-0198, Japan; Institute of Basic Medical Sciences, College of Medicine, National Cheng Kung University, Tainan, 701, Taiwan; School of Pharmaceutical Sciences, University of Shizuoka, Shizuoka, 422-8526, Japan

**Keywords:** Enterohemorrhagic *Escherichia coli*, Shiga toxin, bacteriophage, inhibitor, SOS response, niclosamide, quinoclamine

## Abstract

**Objectives:** Enterohemorrhagic *Escherichia coli* (EHEC) is responsible for the most severe symptoms of *E. coli* infections, including hemorrhagic colitis and hemorrhagic uremic syndrome. Shiga toxin 2 (Stx2) plays a significant role as a major virulence factor. The genes encoding Stx2 locate in lambda-like prophage on the EHEC genome. Consequently, Stx2 is expressed when production of the phage is induced by the SOS response. Antibiotic treatment is not recommended for curing the bacterial infection, because it is associated with severe hemorrhagic uremic syndrome. If Stx2 production is prevented, EHEC pathogenicity significantly decreases, and antibiotics may be available to treat the infection.

**Methods:** We conducted two independent screenings to identify Stx2 production inhibitors for libraries from the RIKEN Natural Product Depository (NPDepo); namely, screening of the Authentic Library, and two-round screening of the Pilot and Analog Libraries.

**Results:** The screening of Authentic Library identified niclosamide as a Stx2 production inhibitor. Besides, two naphthoquinoids were identified after the two-round of screening of the Pilot and Analog Libraries. Niclosamide, and quinoclamine, which has structure shared in the two naphthoquinoids, prevented cell lysis via the phage production and ceased Stx2 production in EHEC. The SOS reporter assay indicated that quinoclamine prevented the SOS response in *E. coli*, whereas niclosamide did not.

**Conclusions:** These findings suggest that quinoclamine inhibited Stx2 production by preventing the SOS response, whereas niclosamide was involved in phage propagation following the SOS response. These compounds can be a potential therapeutic option to treat EHEC infections.

## Introduction

Although many *Escherichia coli* strains are commensal bacteria, some display pathogenicity and are associated with various pathotypes. Among these, enterohemorrhagic *E. coli* (EHEC) infection is the most severe and can result in life-threatening symptoms, especially in elderly people and young children.^1, 2^ As EHEC infections can be caused by food and water contaminations, outbreaks of the infection have often been reported, mainly in America, Europe, and Japan.^3–5^ As many wild and domestic animals are identified as EHEC reservoirs, there is concern that the pathogen can be transmitted through fecal contamination of agricultural products or aquatic environments.^5, 6^ Processed meat and fresh vegetables are listed as risk factors for EHEC infection.^7, 8^ Symptoms of EHEC infection include mild watery diarrhea, hemorrhagic colitis, and, in severe cases, hemorrhagic uremic syndrome, which is associated with a high mortality rate.^1, 2^ However, antibiotic treatment is not recommended to cure EHEC infections, because it has been identified as a high-risk factor for hemorrhagic uremic syndrome.^2, 9, 10^ Therefore, supportive therapy, including hydration and intravenous fluids, is recommended to treat EHEC infections.

Shiga toxin is the major virulence factor of EHEC, but other virulence factors have also been identified.^2, 11^ That is, EHEC, which causes hemorrhagic colitis in humans, is a Shiga toxin-producing *E. coli* (STEC). Shiga toxin is an AB toxin consisting of one A subunit and five B subunits. The subunit B pentamer participates in the interaction with globotriaosylceramide as a receptor on the host cell surface, incorporation into the host cell, and intracellular trafficking.^12^ Subunit A in the delivered protein complex is cleaved by furin, a host protease, to release a part of subunit A from the protein complex.^13^ Consequently, the toxin becomes activated in the cytoplasm. The activated Shiga toxin, an RNA-*N*-glycosidase, removes a specific adenine nucleobase from the 28S rRNA in ribosomes, resulting in inhibition of protein synthesis.^14^ The halt of protein synthesis induces ribotoxic stress and apoptosis, ultimately killing intoxicated cells.^12, 15^ Shiga toxin has been classified into two phylogenetic branches based on biochemical characterization: Shiga toxin 1 (Stx1), which is almost identical to the Shiga toxin from *Shigella dysenteriae*, and Shiga toxin 2 (Stx2), which is not neutralized by polyclonal antisera against Stx1. Most EHEC strains possess multiple toxins, whereas others harbor only one toxin.^16^ Epidemiological studies have shown that EHEC producing Stx2 is associated with more severe symptoms than that not producing Stx2.^11, 17, 18^ *In-vitro* assays showed that Stx2 has a dramatically higher cytotoxicity than Stx1 in human renal microvascular endothelial cells, which are putative target cells for the development of hemorrhagic uremic syndrome.^19^ Furthermore, in a mouse model, Stx2 displayed higher pathogenicity and was significantly associated with hemorrhagic uremic syndrome.^20, 21^

The genes encoding both toxins, *stx1AB* and *stx2AB*, are located in prophages, which are lysogenized into the *E. coli* chromosome. However, since DNA damage does not induce production of the prophage including *stx1AB*, the phage is likely defective.^22^ Conversely, the phage including *stx2AB* is produced by ciprofloxacin or mitomycin C (MMC) treatment.^23, 24^ Although these toxin genes are located at similar loci in the prophage genomes, which are regions for late genes, the transcriptional regulations of these genes are different.^25, 26^ The promoter for *stx1AB* includes Fur box, and the genes are expressed under iron depletion conditions.^27^ Since the prophage including *stx1AB* is not produced, the expressed Stx1 is accumulates in the cells without cell lysis.^28^ In contrast, *stx2AB* genes are expressed during phage production.^29, 30^ The SOS response is activated in *E. coli* cells under certain stresses, such as antibiotic treatment, which is followed by production of the phage including *stx2AB*. Consequently, Stx2 is expressed as a part of late genes during the phage production. Finally, the produced Stx2 is aggressively released through the cell lysis via phage production.^28^ This is another reason for why antibiotics are not recommended to treat EHEC infection.^1, 31^ However, when Stx2, as the most severe virulence factor, is not produced in EHEC, the pathogenicity of the bacterium is dramatically reduced. We thus aimed at identifying chemical compounds that inhibit Stx2 production in EHEC.

## Materials and methods

### Bacterial strains, plasmid, and culture media

The strains and primers used in this study are listed in Tables S1 and S2, respectively. Bacteria were cultured in lysogeny broth (LB) at 37°C with vigorous shaking.^32^ When required, antibiotics and inducers were added at the following concentrations: ampicillin (Ap), 50 µg/mL; kanamycin (Km), 50 µg/mL; tetracycline (Tc), 5 µg/mL; and arabinose, 0.2%. Phage 933W was prepared from EDL933 upon MMC treatment, and it was lysogenized into the MG1655 chromosome.^23^ Lysogenization at the *wrbA* locus and single lysogeny were confirmed via PCR. To construct the MH818 strain for screening, the *gfp* gene from pFPV25.1 was amplified via PCR using the primers EDL-stx2a-GFP and GFP-C-P1, and the Km^R^ cassette from pKD13 was amplified using the primers P1-pKD13 and EDL-stx2b-P2. The amplified fragments were then combined using overlap extension PCR with the primers EDL-stx2a-GFP and EDL-stx2b-P2, and subsequently used for Red recombination with the MG1655 933W/pKD46 strain to replace *stx2AB* with *gfp*-Km^R^.^33^ The Km^R^ cassette in between the FRT sites was then removed by pCP20. To construct pSulA-mCherry as an SOS reporter plasmid, a fragment including the *sulA* promoter, which is regulated by the SOS response, was amplified from the MG1655 chromosomal DNA using the primers 198-28 and 198-29. A fragment for the mCherry gene was amplified from pET-mCherry-LIC with the primers mCherry-N and BII-pG+_MCS, and a fragment of the vector was amplified with the primers CmN-R-BII and CmC-R-E. These three amplified fragments were assembled via SLiCE cloning.^34^

### Chemical libraries from NPDepo

The Authentic, Pilot and Analog Libraries from NPDepo were provided by RIKEN, Japan. The Authentic Library included a curated collection of 80 compounds with extensively documented biological activities. This library contained antibiotics, antitumor and anti-inflammatory drugs, and compounds with protein synthesis inhibitory and corticosteroid-like activities. The list of these compounds is available online.^35^ For the Pilot Library, 25,000 chemical compounds in NPDepo were classified into 400 clusters based on structural similarity. Thus, the compounds in each cluster share the same skeleton or common substructure. One representative compound was selected from each cluster, and the Pilot Library was constructed. To date, the structures of compounds in the Pilot Library have not been disclosed for blind screening. Hit compounds from the Pilot Library were applied for a structural similarity search for 25,000 compounds as the NPDepo library (detail will be published elsewhere). Clusters of analog compounds for each hit compound were assembled as the Analog Library.

### Screening conditions for compounds that inhibit Shiga toxin expression

The Authentic, Pilot, and Analog Libraries provided by NPDepo were used for the screening. The chemical compounds were solubilized in DMSO at concentrations of 1.7 and 8.3 mM. The cultures for screening were conducted in 96-well plates. Briefly, 2 µL of an overnight culture of MH818 was inoculated to 200 µL of fresh LB medium supplemented with 0.5% MMC and 2 µL of the DMSO solution containing chemical compounds (with final concentrations of 17 and 83 µM for the low and high doses, respectively). The culture was incubated at 37 °C with vigorous rotation, and OD_600nm_ and GFP fluorescence intensity were determined every hour for 8 h using a plate reader (SpectraMax M3, Molecular devices).

### Western blotting to determine Shiga toxin levels in EHEC

Overnight culture of EDL933 was inoculated into fresh LB medium with or without 0.5% MMC, niclosamide, and quinoclamine and cultured at 37 °C with vigorous shaking. OD_600nm_ was measured hourly for 8 h, and the cells were harvested via centrifugation for Western blotting. Equal amounts of whole-cell lysates were separated using sodium dodecyl sulfate-polyacrylamide gel electrophoresis. The resolved proteins were transferred onto a PVDF membrane (Millipore) using a semi-dry blotting system (Bio-Rad). Monoclonal antibodies against Stx2A (Hycult Biotech) and homemade OmpA antiserum^36^ were used to detect Stx2A and OmpA, respectively. A horseradish peroxidase-conjugated goat anti-mouse IgG (Abcam) was used as the secondary antibody. Target proteins were visualized using Immobilon Western Chemiluminescent HRP substrate (Millipore).

### SOS reporter assay

An overnight culture of MG1655/pSulA-mCherry was diluted 1:100 with fresh LB medium containing 0.5% MMC, with or without 7 µM niclosamide and 20 µM quinoclamine, and cultured at 37 °C with vigorous shaking. The fluorescence intensity of mCherry was determined every hour for 8 h using a plate reader (SpectraMax M3, Molecular devices).

### Statistical analysis

All experiments were conducted in triplicate and the data are presented as the mean and standard deviation. Student’s t-test was used for statistical analyses. Asterisks in the figures indicate *p*-values with significant differences (\**p*<0.05, \*\**p*<0.001, \*\*\**p*<0.0001).

## Results

### Development of a screening system to identify Stx2 expression inhibitors

In this study, *E. coli* O157:H7 EDL933 was used as the model strain for EHEC.^37^ However, owing to biosafety concerns, a virulence factor-free *E. coli* strain derived from *E. coli* K-12 MG1655 was also constructed for chemical library screening. Prophage 933W, which includes the *stx2AB* genes in EDL933, was activated upon mitomycin C (MMC) treatment, and the isolated phage was lysogenized into the MG1655 chromosome. Furthermore, the genes encoding Stx2 in the lysogenized 933W were replaced with the *gfp* gene. The resulting strain (MH818) was then used for chemical screening. To evaluate the screening system, RecX, a proteinous inhibitor of SOS induction,^38^ was expressed in MH818 (Figure S1). Cell lysis via phage production under MMC treatment was determined using the growth curve, and the expression of GFP was used as a reporter for Stx2 expression. RecX expression indicated that the decrease in OD_600nm_ due to phage production was restored, and GFP expression was significantly reduced. These results indicate that the strain worked well for inhibitor screening.

### Screening of the Authentic Library for Stx2 expression inhibitors

First, the Authentic Library from RIKEN consisting of 80 compounds with previously known biological activities was screened for Stx2 expression inhibitors. Each compound at a concentration of 17 and 82 µM for low and high doses, respectively, was added to the culture with MMC, and the OD_600nm_ and GFP fluorescence intensities were determined every hour for 8 h. Figure 1A shows the relative GFP fluorescence intensity after cells had been cultured with the high dose of chemical compounds, and Figure S2 shows the time course-dependent growth curve and GFP intensity during the culture. The cultures with twelve compounds significantly decreased the GFP intensity to less than 70% at 8 h (Figure 1A). However, because nine of the twelve compounds were well-known antibiotics, three compounds excluding the antibiotics were chosen as hit compounds (Figure 1C). These three compounds were apomorphine hydrochloride hemihydrate (E3), hycanthone (G8), and niclosamide (H5). The dose-dependent effect of these compounds was shown through the lower reduction in GFP expression upon treatment with low doses of the compounds compared to high doses (Figure 1B). Among the three compounds, niclosamide also reduced bacterial cell lysis, indicating its potential to reduce both phage and Stx2 production (Figure S2).

**Figure 1.**
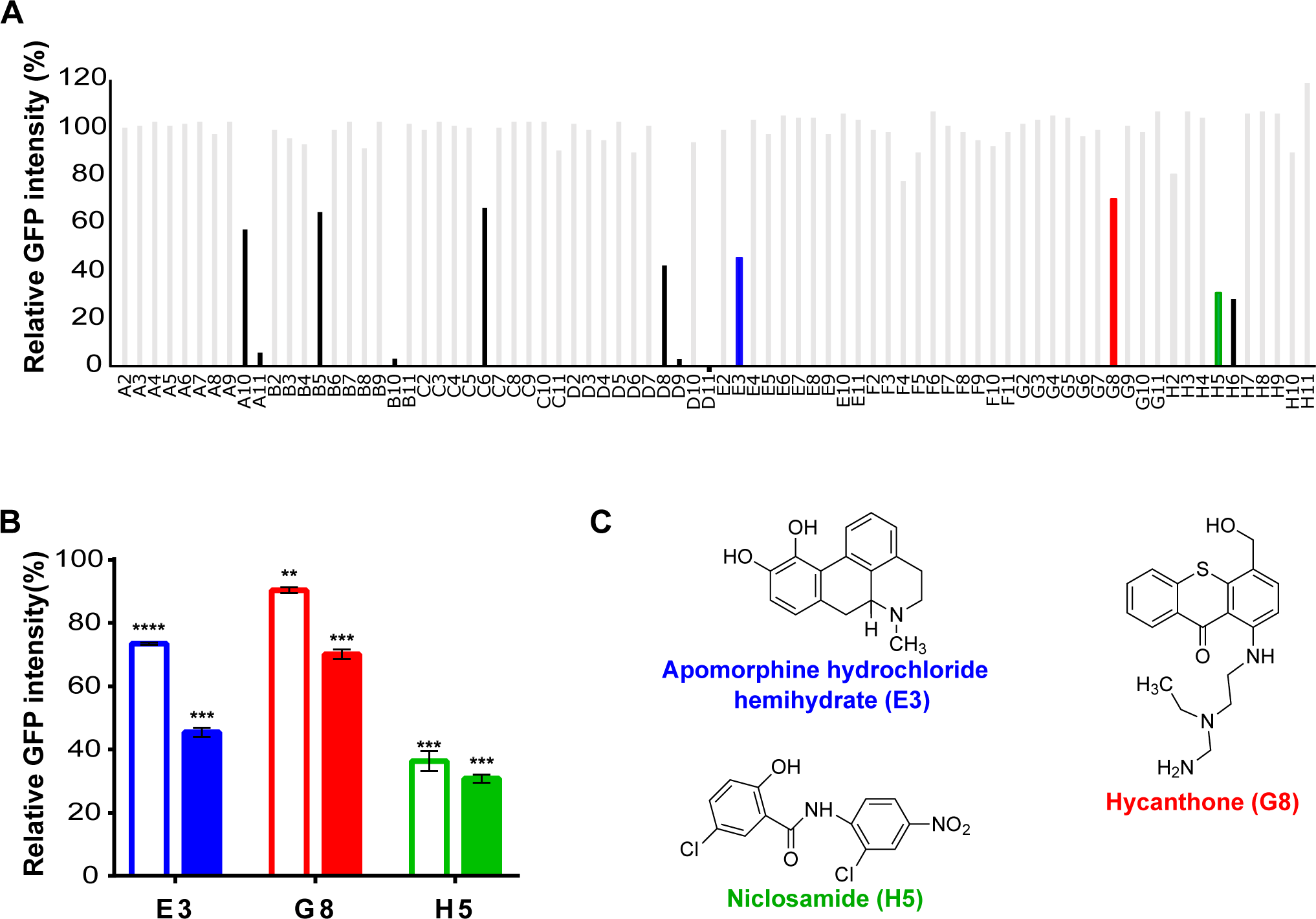
Screening results using the Authentic Library. The Authentic Library contains 80 known bioactive compounds. The screening strain MH818 was cultured with mitomycin C (MMC) and a compound from the Authentic Library. OD_600nm_ and GFP intensity were measured every hour for 8 h. (A) Relative GFP intensity at the 8-h time point with 80 screening compounds from the Authentic Library (83 µM). The experiments were performed in triplicate, and the averages of the results are indicated. Three compounds shown in blue (E3), red (G8), and green (H5) exhibited a significant decrease in GFP expression. Results with known antibiotics are indicated in black, and non-hit compounds are in gray. (B) The hit compounds exhibited a dose-dependent effect in inhibiting GFP expression. Open bar, 17 µM; closed bar, 83 µM. Asterisks indicate a significant difference between the mock and experimental samples as determined using Student’s t-test (\**p*<0.05, \*\**p*<0.001, \*\*\**p*<0.0001). (C) Chemical structures of the hit compounds: E3, apomorphine hydrochloride hemihydrate (blue); G8, hycanthone (red); H5, niclosamide (green).

To demonstrate this in a pathogenic strain, EDL933 was cultured with or without MMC and niclosamide, and the growth curve and expression of Stx2 were determined (Figure 2). The growth curve showed a reduction in OD_600nm_ when only MMC was added to the EDL933 culture, suggesting that the phage and Stx2 were produced by MMC treatment. However, when both MMC and niclosamide were added, the reduction in OD_600nm_ was significantly slower than that upon treatment with MMC only. The amount of Stx2 was dramatically reduced when the concentration of niclosamide exceeded 6 µM (Figure 2B). These results demonstrate that niclosamide reduces Stx2 expression in EHEC and support the utility of our screening system for identification of compounds that inhibit Stx2 expression.

**Figure 2.**
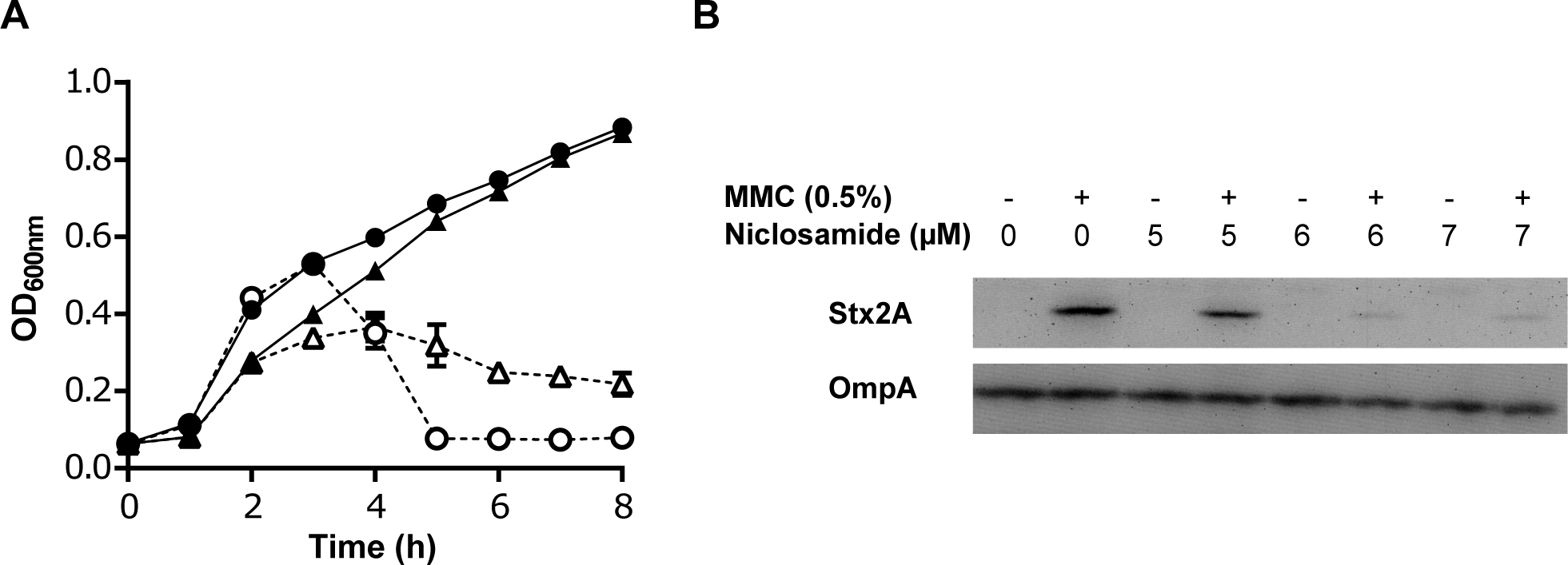
Niclosamide reduces Shiga toxin expression in EHEC. (A) Growth curve of EDL933 treated with niclosamide. EDL933 was cultured with or without MMC (dashed and solid lines, respectively), and with or without 7 µM niclosamide (triangles and circles, respectively). (B) Western blotting for Shiga toxin. Cells in the cultures described in panel A were harvested at the 8-h time point, and the expressions of Stx2 and OmpA were visualized.

### Screening of the Natural Products Depository (NPDepo) Library

Because our screening system could identify inhibitors of Stx2 expression as described above, we applied it to screen a library containing approximately 25,000 natural compounds, i. e., the RIKEN Natural Products Depository (NPDepo), Japan. However, screening for a large number of compounds is time-consuming, labor-intensive, and the hit frequency is very low. To address this, we employed a two-round strategy for the screening process. First, the compounds in the library were classified into 400 clusters based on their structural similarity; therefore, the compounds in each cluster shared part of their structures. One compound that typically symbolized each cluster was chosen to represent that cluster, and the 400 representative compounds were assembled as Pilot Library. Consequently, the structures of the compounds in the Pilot Library were different from one another, so the library has richness in structural diversity. Screening of the Pilot Library was performed under the same conditions as those used for screening of the Authentic Library. Screening of the Pilot Library as the first round indicated that 17 compounds significantly decreased GFP expression at the high dose, and this effect was dose-dependent (Figure 3 and S3). Among them, 11 compounds, excluding six as known antibiotics, were selected as potential hit compounds. Then, compounds structurally similar to these 11 compounds were selected from the original NPDepo Library, and 11 clusters with analog compounds were constructed. In total, 160 compounds were included in the 11 clusters and used as Analog Library for the second round of screening (Figure 4). Although NPD1B7 and NPD1C4 in the Analog Library showed the lowest GFP expression levels at 8 h, they also hindered bacterial growth. In contrast, NPD1G3 and NPD2G11 did not reduce OD_600nm_ at the late culture stage and showed relatively lower GFP expression than the other compounds. The cultures containing NPD1G3 and NPD2G11 appeared to exhibit normal growth without bacterial lysis, suggesting that the productions of phage and Stx2 were prevented. Therefore, we focused on these compounds in subsequent experiments.

**Figure 3.**
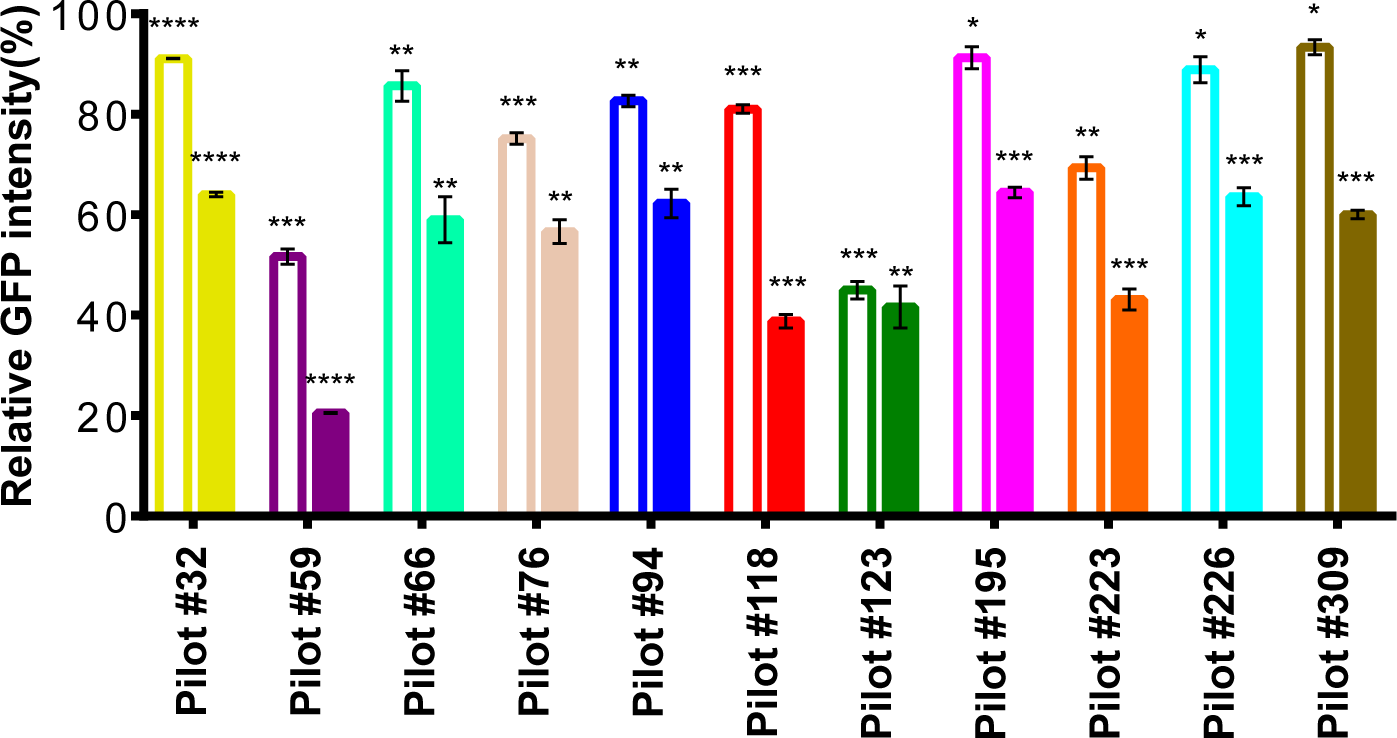
Screening results using the Pilot Library. Four hundred compounds in the Pilot Library were applied for screening (Figure S3), wherein 11 compounds displayed a significant reduction in GFP expression. Open bar, 17 µM; closed bar, 83 µM. Asterisks indicate a significant difference between the mock and experimental samples as determined using Student’s t-test (\**p*<0.05), \*\**p*<0.001), \*\*\**p*<0.0001).

**Figure 4.**
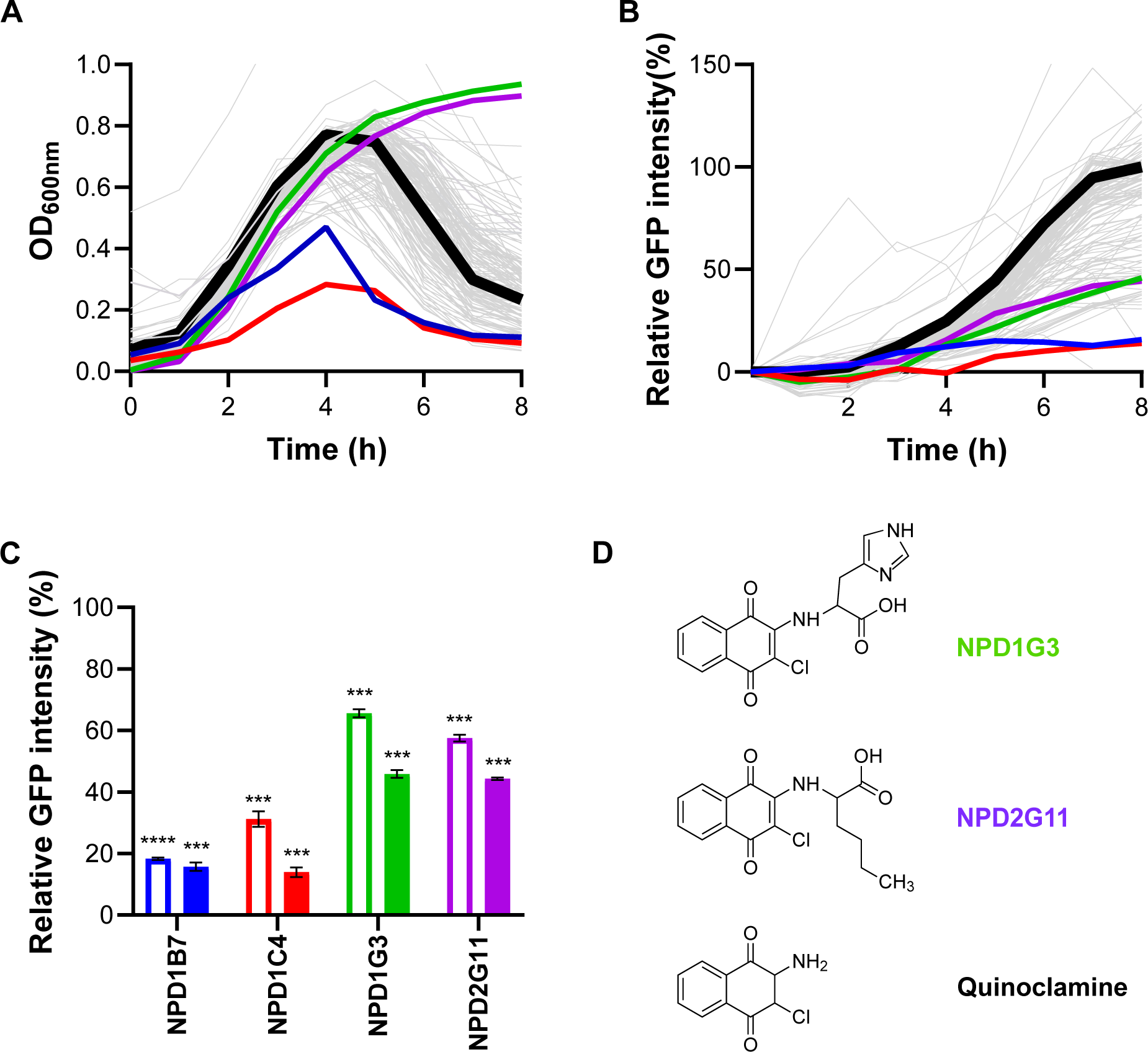
Screening results using the Analog Library. The Analog Library, which contains 160 compounds, was applied for the screening. The time course changes in OD_600nm_ and GFP intensity are indicated in panels (A) and (B), respectively. The results of the mock samples are indicated by the thick black line. Non-hit compounds are shown by gray lines. The compounds described in the text are indicated in colors: blue, NPD1B7; red NPD1C4; green, NPD1G3; and purple, NPD2G11. (C) The hit compounds exhibited a dose-dependent effect in inhibiting GFP expression (open bar, 17 µM; closed bar, 83 µM). Asterisks indicate a significant difference between the mock and experimental samples as determined using Student’s t-test (\**p*<0.05, \*\**p*<0.001, \*\*\**p*<0.0001). (D) Chemical structures of NPD1G3, NPD2G11, and quinoclamine. Following the NPDepo policy, the structures for NPD1B7 and NPD1C4 were not disclosed (see Materials and Methods).

NPD1G3 and NPD2G11 belonged to the same analog cluster, which indicates that they have similarities in their structures. The compounds in the analog cluster that included NPD1G3 and NPD2G11 are illustrated in Figure S4. Only NPD1G3 and NPD2G11 showed normal growth in the analog cluster, whereas the other analogs reduced OD_600nm_ (Figure S4). The molecular structures of NPD1G3 and NPD2G11 share the 2-amino-3-chloro-1,4-naphthoquinone moiety, although the other compounds were not chlorinated at position 3 (Figure S4). The structure-function relationship of the compounds led us to hypothesize that the shared structure of NPD1G3 and NPD2G11 plays a role in inhibiting bacterial cell lysis and GFP expression. Quinoclamine, shown in Figure 4D as 2-amino-3-chloro-1,4-naphthoquinone, was then used to determine whether it prevents Stx2 production in pathogenic EHEC (Figure 5). Supplementation of quinoclamine to the culture without MMC did not impact growth. However, increasing the concentration of quinoclamine in the culture with MMC restored bacterial growth; the culture containing 8 µM quinoclamine grew normally, even when MMC was supplemented. Moreover, Stx2 expression in EDL933 under MMC treatment dramatically decreased when quinoclamine was added (Figure 5B). These results clearly demonstrate that quinoclamine prevented Stx2 production in the pathogenic EHEC strain.

**Figure 5.**
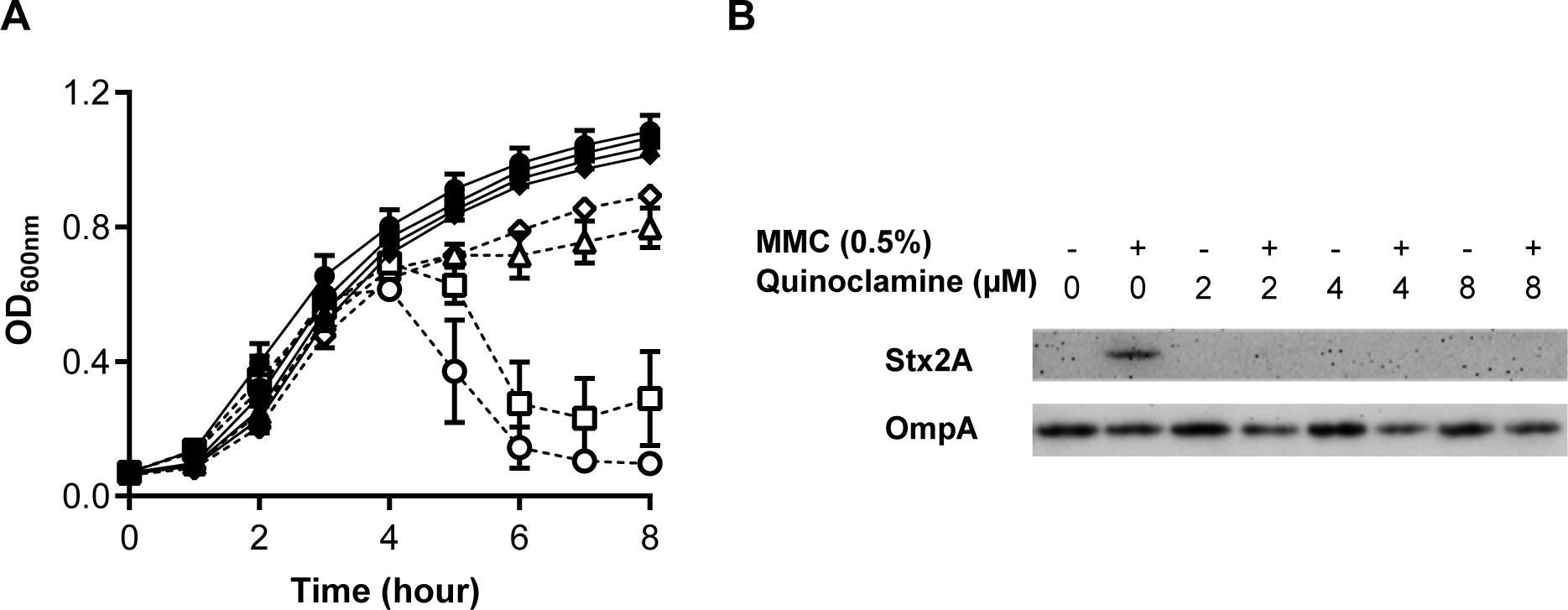
Quinoclamine reduces Shiga toxin expression in EHEC. (A) Growth curve of EDL933 cultured with quinoclamine. EDL933 was cultured with or without MMC (dashed and solid lines, respectively) and different concentrations of quinoclamine (circle, 0 µM; square, 2 µM; triangle, 4 µM; diamond, 8 µM). (B) Western blotting for Shiga toxin. Cells in the cultures described in panel A were harvested at the 8-h time point, and the expressions of Stx2 and OmpA were visualized.

### Quinoclamine reduces the SOS response, while niclosamide did not

The production of the phage 933W, including *stx2AB*, is regulated by the SOS response in *E. coli* cells, and genes encoded in the phage genome are sequentially expressed under SOS induction. To investigate whether the inhibitors found in this study were involved in the SOS response or phage propagation, the level of SOS induction was determined for niclosamide and quinoclamine (Figure 6). A reporter strain for the SOS response, which does not contain the lysogenized phage 933W, was cultured with or without these compounds. Although niclosamide did not affect the SOS response, quinoclamine restored SOS induction to the basal level. These results suggest that quinoclamine prevents the SOS response, whereas niclosamide hinders phage propagation.

**Figure 6.**
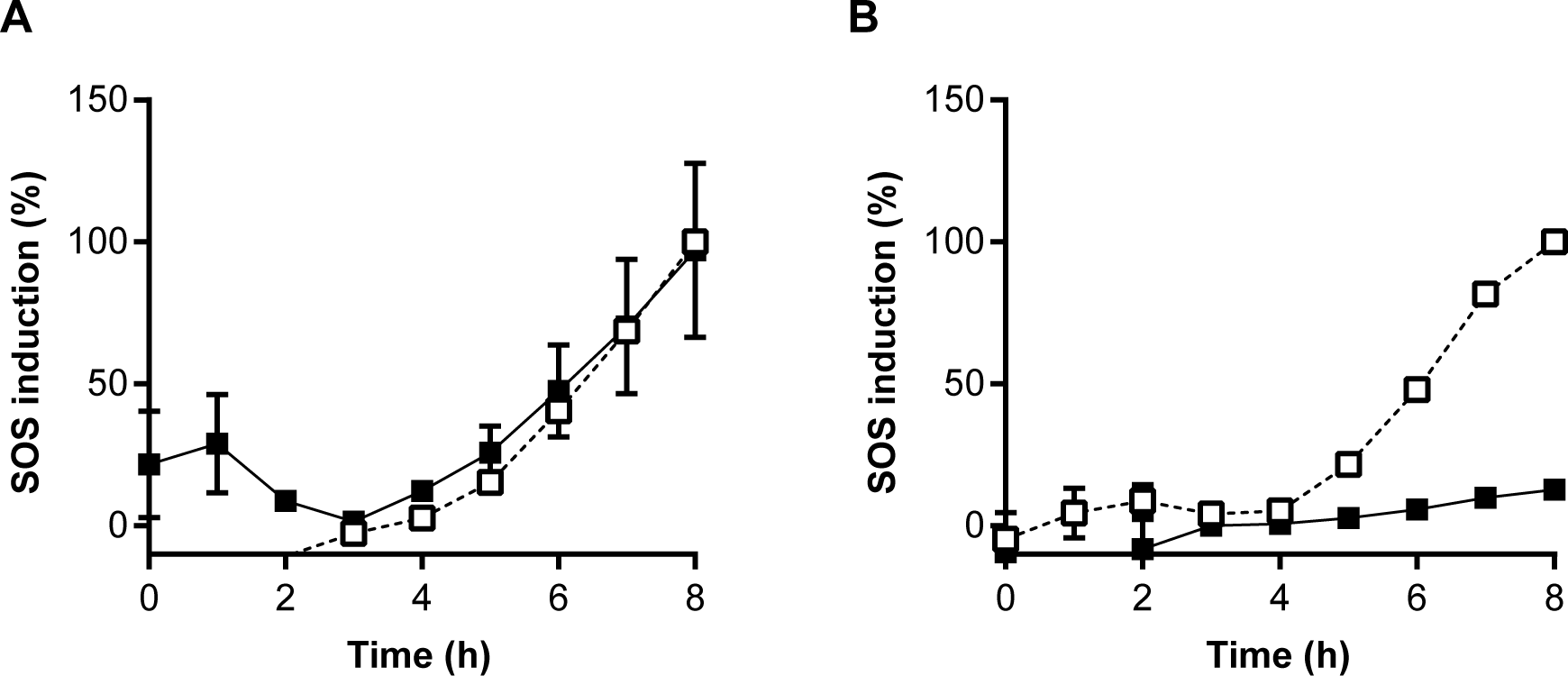
SOS reporter assay with quinoclamine and niclosamide. An SOS reporter strain (P*_sulA_*-mCherry) was cultured with or without MMC, niclosamide and quinoclamine, and the fluorescence intensity of mCherry was determined. MG1655/pSulA-mCherry was cultured in LB containing 0.5% MMC (dashed line). Inhibitors were added (panel A, 7 µM niclosamide; panel B, 20 µM quinoclamine) and indicated by a solid line.

## Discussion

In this study, inhibitors for Shiga toxin production were identified through two independent screenings. Namely, niclosamide was identified from the Authentic Library, and quinoclamine was identified through two-round screening of the Pilot Library and the Analog Library (Figure S5). These compounds significantly reduced Shiga toxin expression in EDL933 as a pathogenic EHEC strain under MMC treatment, allowing EHEC growth without cell lysis via phage production. This suggests that the pathogenicity of EHEC was substantially reduced. The two-round screening of the NPDepo, which includes approximately 25,000 natural compounds, demonstrated a high hit rate and efficiency. Screening for all of the compounds would have been considerably less efficient in identifying hit compounds and would have required considerably more time and labor.

The second round of screening for the Analog Library, which is consisted of 11 analog clusters, revealed three patterns. The first pattern indicated that only selected representative compounds within each cluster significantly reduced GFP expression at 8 h. Five clusters, including NPD1C4, displayed this pattern. In this case, selection of representative compounds is important. It implies that hit compounds may still exist in clusters that have not yet been examined. The second pattern indicated that the compounds within each cluster exhibited various efficacies. Three clusters, including NPD1G3 and NPD2G11, showed this pattern. In this case, both of the common backbone structure and side chains may play a role in the function. The third pattern indicated that all analogs showed a similar reduction in GFP expression. Three clusters, including NPD1B7, exhibited this pattern, in which the shared backbone structure may be important for functioning. In the latter two patterns, the two-round screening strategy worked well for the screening efficiency, as GFP expression was reduced stepwise from round to round. Thus, only 560 of 25,000 compounds were tested, and quinoclamine was effectively identified.

Niclosamide is included in the WHO Model List of Essential Medicines as an intestinal anthelmintic for cestode infections.^39^ This compound shows multiple biological activities, including anti-inflammatory, antimicrobial, anti-tumor, and bronchodilatory activities.^40, 41^ Moreover, it has recently been studied in treating COVID-19.^42–44^ In the past decade, niclosamide has been shown to have antibacterial activity against Gram-positive bacteria, including *Staphylococcus aureus* and *Clostridium difficile*. However, whether niclosamide exhibits antibiotic activity against Gram-negative bacteria remains unknown, although it enhances the efficacy of colistin in Gram-negative bacteria.^45^ In the first screening in this study, 83 µM niclosamide allowed *E. coli* growth, suggesting that it did not exhibit antibiotic activity (Figure S2). Furthermore, niclosamide did not suppress the SOS response in *E coli* (Figure 6). Taken together, these results suggest that niclosamide prevents phage propagation after the SOS response. Since multiple genes regulate lambdoid phage production, further studies are required to identify the targets of niclosamide.

Quinoclamine derivatives were effectively isolated as inhibitors of Shiga toxin production through our two-round screening strategy. In the second round of screening, the inhibitory activities of naphthoquinoids were compared per cluster. Only NPD1G3 and NPD2G11, which were chlorinated at position 3, prevented the decrease in OD_600nm_ (Figure S4). Moreover, comparing these two compounds led us to propose that modifying the amino group at position 2 is not important to their biological function. Indeed, quinoclamine inhibited Shiga toxin production even without any amino group modification (Figure 5). Thus, the structure-function analysis suggested flexibility of the amino group. Quinoclamine or its derivatives can be a potential therapeutic option for EHEC infection. To date, quinoclamine is used as a herbicide that inhibits photosynthesis in plants.^46, 47^ Recently, it has been used as an inhibitor for NF-κB.^48^ In addition, its derivatives showed antiplatelet, anti-inflammatory, antiallergy, and inhibitory effects on MEK1.^49, 50^ Therefore, medical applications of these compounds are anticipated. In this study, quinoclamine prevented Shiga toxin production through inhibiting the SOS response. To the best of our knowledge, this is the first report on the biological activity of quinoclamine against bacteria. However, since quinoclamine treatment up to 100 µM did not affect *E. coli* growth, it might not have antibiotic activity (data not shown).

The SOS response regulates the expression of genes involved in DNA repair, mutagenesis, and the cell cycle. Lambdoid phages utilize the SOS response to switch from lysogeny to the lytic cycle; therefor, the SOS response triggers Shiga toxin expression. Consequently, quinoclamine, which inhibits the SOS response, was identified by screening for inhibitors of Shiga toxin expression. The SOS response increases the frequency of obtaining a mutation for antibiotic resistance and also promotes horizontal gene transfer, which facilitates the spread of antibiotic resistance genes.^51, 52^ The increase in the prevalence of antibiotic-resistant bacteria is a major global health issue, and the lack of novel antibiotics exacerbates this problem. Therefore, developing an antimutagenic agent that can decrease the frequency of obtaining antibiotic resistance is essential, and several inhibitors of the SOS response have been studied for this purpose.^53^ However, their application has not yet been established. Quinoclamine is a potential candidate for reducing mutations by inhibiting the SOS response. Moreover, because the SOS response is conserved in most bacteria, quinoclamine might also be applicable to other bacteria.^54^ The molecular mechanism through which quinoclamine inhibits SOS response is currently being investigated.

## Acknowledgments

We thank the NBRP (NIG, Japan) for *E. coli* strains and plasmids; WPZ for technical supports.

## Transparency declarations

Conflict of interest: none to declare.

## Supplemental Material

Figure S1. De**m**onstration of screening for Shiga toxin-expression inhibitor.

Figure S2. Time course of screening of the Authentic library.

Figure S3. Screening result of the Pilot library.

Figure S4. Time course of the second-round screening for analogues of NPD1G3 and NPD2G11.

Figure S5. Summary of the screenings.

Table S1. Strain and plasmid used in this study.

Table S2. Primers used in this study.

